# crAssphage abundance and genomic selective pressure correlate with altered bacterial abundance in the fecal microbiota of South African mother-infant dyads

**DOI:** 10.1101/582015

**Authors:** Bryan P. Brown, Jerome Wendoh, Denis Chopera, Enock Havyarimana, Shameem Jaumdally, Donald D. Nyangahu, Clive M. Gray, Darren P. Martin, Arvind Varsani, Heather B. Jaspan

## Abstract

crAssphages are a class of bacteriophages that are highly abundant in the human gastrointestinal tract. Accordingly, crAssphage genomes have been identified in most human fecal viral metagenome studies. However, we currently have an incomplete understanding of factors impacting the transmission frequencies of these phages between mothers and infants, and the evolutionary pressures associated with such transmissions. Here, we use metagenome sequencing of stool-associated virus-like particles to identify the prevalence of crAssphage across ten South African mother-infant dyads that are discordant for HIV infection. We report the identification of a complete 97kb crAssphage genome, parts of which are detected at variable levels across each mother-infant dyad. We observed average nucleotide sequence identities of >99% for crAssphages from related mother-infant pairs but ∼97% identities between crAssphages from unrelated mothers and infants: a finding strongly suggestive of vertical mother to infant transmission. We further analyzed patterns of nucleotide diversity across the crAssphage sequences described here, identifying particularly elevated positive selection in RNA polymerase and phage tail protein encoding genes, which we validated against a crAssphage genome from previous studies. Using 16S rRNA gene sequencing, we found that the relative abundances of *Bacteroides thetaiotaomicron* and *Parabacteroides merdae* (Order: Bacteroidales) were differentially correlated with crAssphage abundance. Together, our results reveal that crAssphages may be vertically transmitted from mothers to their infants and that hotspots of selection within crAssphage RNA polymerase and phage tail protein encoding genes are potentially mediated by interactions between crAssphages and their bacterial partners.

**Importance:** crAssphages are an ubiquitous member of the human gut microbiome and modulate interactions with key bacterial associates within the order Bacteroidales. However, the role of this interaction in the genomic evolution of crAssphage remains unclear. Across a longitudinally sampled cohort of ten South African mother-infant dyads, we use metagenome sequencing of the fecal virome and 16S rRNA gene sequencing of the fecal bacterial microbiota to elucidate the ecological and evolutionary dynamics of these interactions. Here, we demonstrate elevated levels of crAssphage average nucleotide identity between related mother-infant dyads as compared to unrelated individuals, suggesting vertical transmission. We report strong positive selection in crAssphage RNA polymerase and phage tail protein genes. Finally, we demonstrate that crAssphage abundance is linearly correlated (P < 0.014) with the abundance of two bacterial taxa, *Bacteroides thetaiotaomicron* and *Parabacteroides merdae.* These results suggest that phage-bacterial interactions may help shape ecological and evolutionary dynamics in the gut.

## Introduction

The human gut virome is dominated by bacteriophages (1-3). In the last decade, a bacteriophage species’ ∼97kb circular DNA genome, commonly referred to as crAssphage (from <u>cr</u>oss <u>ass</u>embly of 12 fecal metagenomes), was identified and found to be the most abundant virus in the human gut (2). crAssphage-like contigs have been *de novo* assembled from fecal samples from various regions of the world and across health and disease states, emphasizing its high prevalence among human enteric microbiota. Among all the currently published complete and partial crAssphage genomes, irrespective of whether they have been isolated from environmental samples, humans or primates (4-6), there exists >90% sequence similarity between homologous genome regions (2, 7-9). Though the evolutionary relationships of crAssphages to other phage families remains unresolved (some evidence points toward a distant relationship to phages in the family *Podoviridae* (10)), comparative genomic analyses of human-associated crAssphage taxa have partitioned them into four candidate subfamilies composed of ten putative genera.

Despite the abundance of crAssphage-like contigs detected in human fecal microbiota, little is known about the lifestyle or selective pressures acting on crAssphage genomes. Based on analysis of CRISPR spacers and other genomic features, it was speculated that crAssphages prey upon members of *Bacteroides* (2). Intuitively, this fits well with the Bacteroidetes-dominated community profile of human gut microbiota. Shkoporov et al. (11) recently confirmed that members of the crAssphage group stably infect *Bacteroides intestinalis* and are able to maintain long-term persistence *in vivo*, though the mechanisms underlying this relationship are unknown. Furthermore, after long-term (23 days) interaction experiments between a crAssphage isolate (crAss001) and *B. intestinalis*, approximately half of isolated *B. intestinalis* colonies demonstrated complete resistance to the phage, indicating rapid evolution of bacterial interaction factors (11). However, it remains unclear how this relationship impacts selective pressures acting across the crAssphage genome.

Our knowledge of bacterial-phage coevolution in the human gut is largely unexplored (12, 13). Antagonistic interactions between bacteria and phages are crucial determinants of genomic evolution for both partners, and several studies have described these trends across diverse systems and environments (14-17). In the gut environment, Minot et al. (12) described rapid nucleotide substitution rates (>10^−5^ per nucleotide per day) in the lytic phage family *Microviridae*. With the establishment of a bacterial host for crAssphage taxa, evolutionary studies of selection in these host cells are likely to reveal some of the genomic consequences of bacterial-phage relationships in the gut.

Here, we report the identification of a complete genome sequence of a crAssphage variant (M186D4) from a South African adult stool sample. To our knowledge, this is the first complete genome obtained from a single individual. Metagenomic sequencing of the viral microbiota of ten mother-infant dyads, that were differentially infected/exposed to HIV, yielded similar levels of crAssphage genomic coverage between mothers and related infants at one week postpartum, potentially indicating comparable crAssphage abundance between related mothers and infants and, therefore, possible vertical transmission. Across the genome of isolate M186D4, we identify selective “hot spots” of nucleotide variation including phage tail protein genes and, to a lesser extent, the RNA polymerase genes. These results were validated across the first published crAssphage genome, illustrating selective pressures acting on human associated crAssphages that potentially arise as a consequence of antagonistic interactions with bacterial consortia in the gastrointestinal tract.

## Materials and Methods

### Sample Collection and virus-like particle isolation

Stools were collected from 10 infants and their mothers from 4 days to 15 weeks post vaginal delivery (Table 2) at a periurban clinic in Cape Town, South Africa. Stools were transported on ice and stored at −80°C until nucleic acid extraction. Samples were defrosted and approximately 0.5g of fecal sample was homogenized in 20ml SM buffer as previously described (18) and centrifuged at 10,000 × g for 10 min. The resulting supernatant was filtered sequentially through a 0.45µm and 0.2µm syringe filter. Filtered supernatants were incubated with lysozyme and Turbo DNAse (Thermo Fisher Scientific, USA) at 37°C for 1 hour to degrade nucleic acids not enclosed in virus-like particles.

### Viral nucleic acid extraction and sequencing

Viral nucleic acid was extracted from 200µl of the filtrate using the High Pure viral nucleic acid kit (Roche Diagnostics, USA) following the standard protocol. Circular viral DNA was amplified using rolling circle amplification (RCA) with Illustra TempliPhi 100 amplification kit (GE Healthcare, USA). The RCA products were used to prepare a 350 bp insert DNA library for each sample following the manufacturer’s standard protocol. Shotgun sequencing was performed on an Illumina HiSeq 2500 platform using 150bp PE chemistry by Novogene (Hong Kong).

### Bacterial genomic DNA (gDNA) extraction and sequencing

Bacterial gDNA was extracted from the same stool samples as described above. Stools were incubated with 6µl mutanolysin (25kU/ml), 50µl lysozyme (450kU/ml), and 3µl lysostaphin (4kU), and incubated at 37°C for 60 minutes with shaking. gDNA was then extracted using the MoBio Powersoil kit following the manufacturer’s instructions. Libraries of the V6 region of 16S rDNA were prepared and generated as described previously (19). Individual libraries were purified using the QIAquick 96 PCR purification kit, quantitated with the Quant-iT dsDNA Broad Range assay, and pooled in equimolar quantities. Pooled libraries were visualized on an agarose gel, excised, and purified using the QIAquick Gel Extraction kit. Library QC and sequencing was performed by the Canadian Centre for Applied Genomics on an Illumina HiSeq 2000 using a 100bp PE approach.

### Viral metagenome assembly and annotation

Raw sequencing reads were trimmed using Trimmomatic v0.36 (20) and then *de novo* assembled using spades v 3.12 (21). All *de novo* assembled contigs of >500 nucleotides (nt) were analyzed using a BLASTx search (22) against a local viral RefSeq protein database compiled from GenBank. An approximate ∼97kb circular (identified by terminal redundancy) contig was identified that had similarities to the first published crAssphage genome (2). Open reading frames (ORFs) were identified using Glimmer (23, 24) and annotated using an in-house reference protein database compiled from NCBI’s GenBank resource.

### Bacterial 16S sequence processing

Forward and reverse indices and marker gene primers were removed using cutadapt (25). Trimmed reads were then quality filtered, dereplicated into amplicon sequence variants (ASVs), and taxonomically classified using the dada2 package (26). Taxonomic classification of ASVs was performed using the Ribosomal Database Project’s Naïve Bayesian classifier (27) against training set 16. Resulting sequence tables were imported into the Phyloseq (28) framework for further processing and analysis. ASV and sample filtering parameters were estimated and applied using custom R functions available at https://github.com/itsmisterbrown/microfiltR. A full vignette detailing our filtering strategy is available at the same location.

### Compositional transformations and bacterial 16S sequence analysis

Filtered datasets were transformed into centered log ratio (CLR) coordinates and subset to taxa within Bacteroidales. Pearson correlations were performed between the CLR transformed abundance of each ASV and the genomic coverage of crAssphage for that sample. Taxa with absolute Pearson’s *r* > 0.4 were included in downstream analyses.

To reduce compositional effects and infer relationships between crAssphage abundance and bacterial abundance, we generated isometric log ratio (ILR) balances of bacterial taxa that were identified as significantly different between samples with high and low crAssphage abundance. Wilcoxon rank sum tests were performed on the CLR abundance of each taxon to determine significance. The ILR transform is a compositional data analytical method that reduces compositional effects by translating relative abundance datasets into orthonormal coordinates suitable for standard statistical analyses (29). Balances enable highly interpretable results of ILR coordinates by using selected sets of taxa in the log ratios. ILR balances were generated using the following equation:

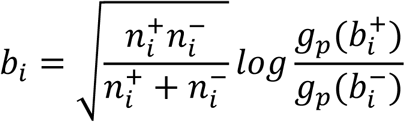

where 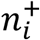 and 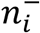 indicate the set of taxa involved the numerator and denominator of that balance (b_i_), respectively, and 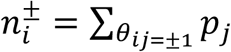 provides the weight integration (30). The first part of the listed balance equation, 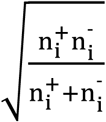, acts as a scaling factor that normalizes each balance to unit length, regardless of weighting scheme. The weighted geometric mean (30) of the taxa associated with a given balance is represented as 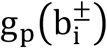 and can be formalized as:

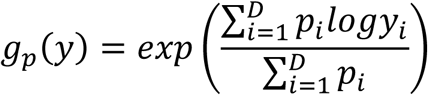

Here, p_i_ represents the weight assigned to a taxon_*i*_, which is either uniform or reflective of its abundance across the dataset. When uniform weights are applied, the geometric mean and isometric log ratio equations default to the original form (29, 31).

### Variant analysis and statistics

Trimmed paired end reads were mapped to the closed M186D4 genome using the BWA with default parameters (32). Samtools mpileup utility (33) was used to calculated the per-nucleotide read coverage and variant frequency. Variants were detected and filtered using VarScan (34). Variants were only considered authentic if there were, at minimum, five high quality reads supporting the variant, which was the lowest threshold for significance in our dataset utilizing an α value of 0.05. P values were calculated using a Fisher’s Exact test on read counts supporting the reference and variant alleles (34).

Functional effects of identified variants were predicted using SnpEff (35) against a custom annotation database generated as part of this study and annotated as nonsynonymous (*N*_*d*_) or synonymous (*S*_*d*_) substitutions. The total number of nonsynonymous (N) and synonymous (S) sites for each protein coding sequence was calculated using SnpGenie (36). The numbers of nonsynonymous (*d*_*N*_) and synonymous (*d*_*S*_) substitutions per site were estimated using the Jukes-Cantor formula:

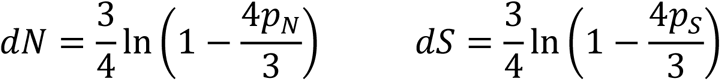

where *p*_*N*_ and *p*_*S*_ indicate the proportions of nonsynonymous and synonymous substitutions, respectively, and can be estimated by 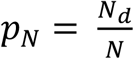 and 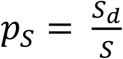.

Genome wide average nucleotide identity (ANI) calculations were performed as described previously (37). Briefly, a sliding window algorithm was used to assess nucleotide identity in 1000bp fragments. Alignments were required to span 200bp and have a minimum identity of 70%. For incomplete genomes generated in this study, trimmed reads were recruited to the genome of isolate M186D4 using Bowtie2 (38) with “very sensitive local” parameters (not requiring end-to-end read alignment). Reads that aligned to genome M186D4 were then used for genome assembly with the spades assembler (39). Assembled contigs were used for ANI calculations for samples without closed genomes. P values for ANI comparisons were calculated from Wilcoxon rank sum tests due to variations in sample sizes between groups. Downstream statistical analysis and visualization was conducted using the base R framework and the ggplot2 package (40).

### Validation of results against a previously published crAssphage genome

The vast majority of publicly available crAssphage genomes have been assembled from pooled samples of many individuals (2, 3, 8, 11), rendering those datasets mostly unsuitable for variant analysis. However, the individual datasets from which the original crAssphage genome sequence was cross assembled (3) were primarily composed of related mother-infant pairs with low interpersonal viral diversity, thus representing the most attractive option for cross validation. Raw nucleotide sequences were download from the NCBI Sequence Read Archive from Study SRP002523. The genome assembly and raw reads from Run SRR073438 were selected for variant analysis. Run SRR073438 was pooled DNA isolated from virus-like particles from a mother and twin pair, as well as an additional maternal sample. Genes and annotations were transferred from the associated published genome (GenBank accession #BK010471) using Prokka (41). Quality filtered reads from each run were aligned to the genome from which they were assembled using BWA (32). For run SRR073438, an α value of 0.05 was used to set read thresholds for variant significance because the vast majority of reads were isolated from a mother and related twins and the broader interpersonal diversity across the dataset was low (3).

### Data availability

The annotated genome sequence for crAssphage M186D4 is available under GenBank accession number MK238400. Partial crAssphage genome sequences are available under NCBI BioProject PRJNA526942. The data, functions, and R script required to reproduce the 16S analyses used in this study are available at https://github.com/itsmisterbrown/crAssphage_M186D4_analyses

## Results

### Metagenome sequencing and assembly

From a 24 year old, HIV-infected female’s stool sample, a closed circular genome of 97,757 kb (Figure 1) was identified as having high similarity (96.6% ANI; Figure 2A) to the first identified crAssphage (GenBank accession #BK010471) (2). The participant was one week postpartum, had a CD4 count of 265 cells/mm^3^, and had initiated antiretroviral therapy during pregnancy. The genome was fully closed with 451-fold mean coverage from 140,984 150bp PE reads (Table 1, Figure 3B). We identified and annotated 82 ORFs and protein coding sequences. Of the 10 candidate genera proposed by Guerin et al (9), crAssphage M186D4 belonged in candidate genus I, with a genome-wide GC content of 29%.

**Table 1.**
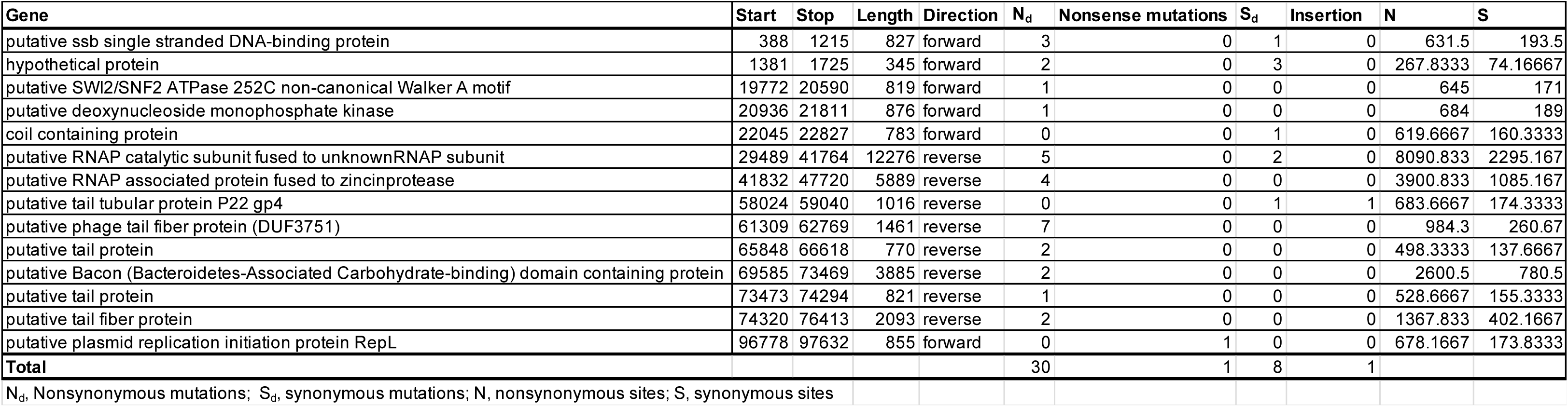
Functional annotation of variants in crAssphage M186D4.

**Figure 1.**
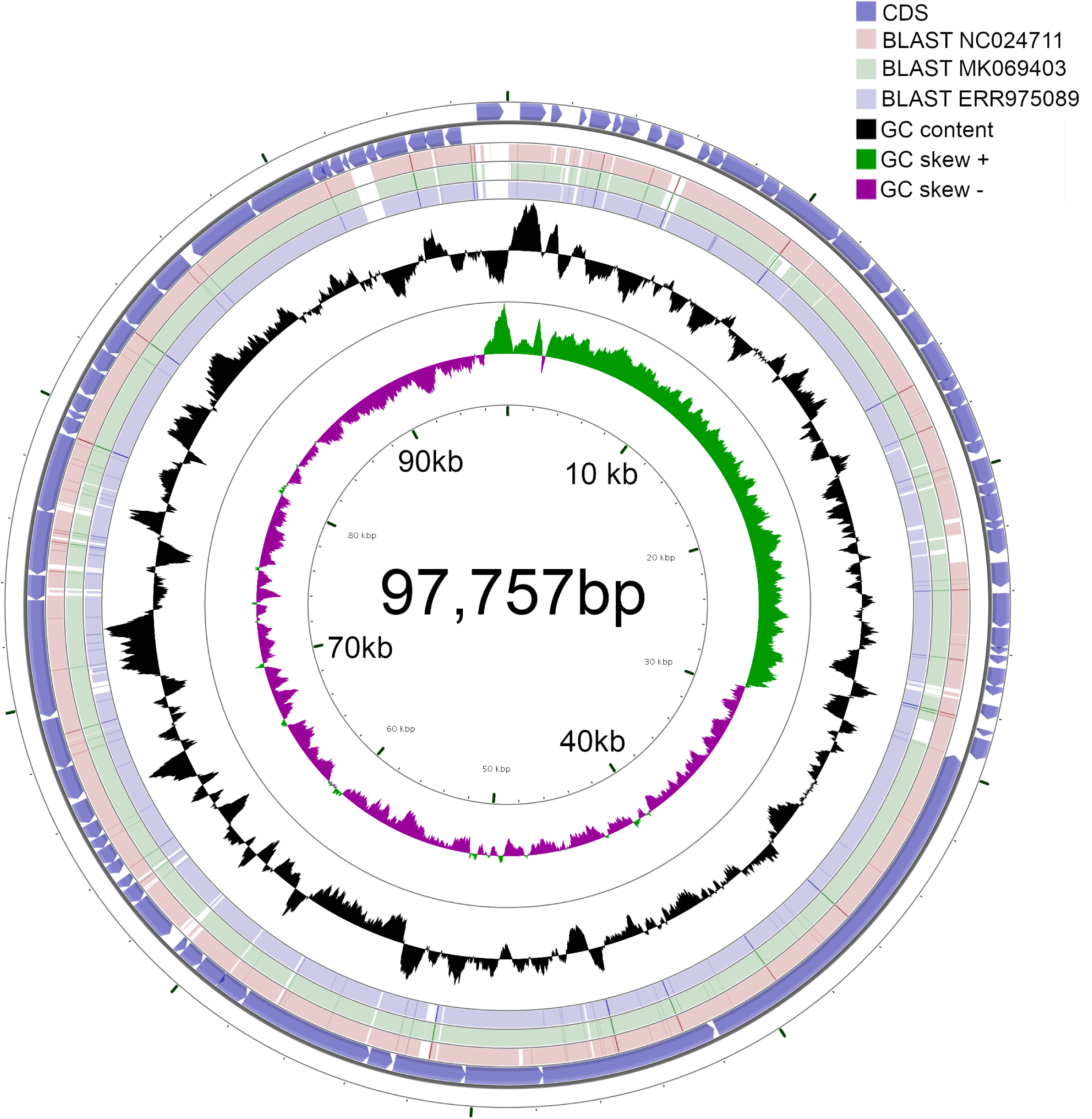
The 97,757bp genome of crAssphage M186D4. CDS are located on the outermost circle and are colored blue. BLAST alignments of isolate M186D4 to three additional crAssphage genomes are colored red green and blue and their GenBank accession number is listed in the legend. GC content is displayed in black. GC skew is displayed in purple and green.

**Figure 2.**
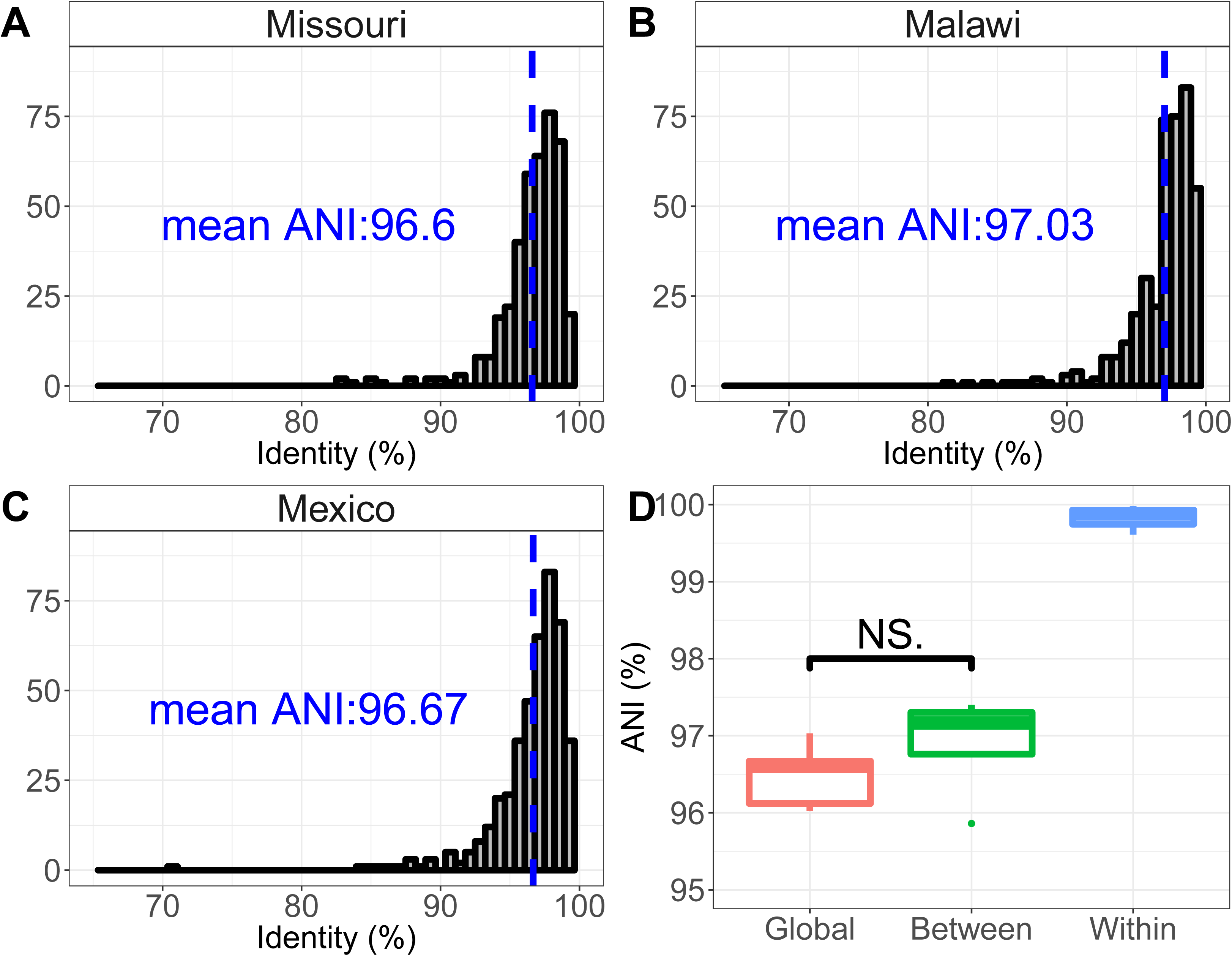
Average nucleotide identity of crAssphage genomes is higher between related mother-infant dyads that between unrelated individuals. **A**. Histogram of the average nucleotide identity of 1,000bp fragments of crAssphage genomes from Missouri (2, 3) and this study. **B**. Histogram of the average nucleotide identity of 1,000bp fragments of crAssphage genomes from Malawi (42) and this study. **C**. Histogram of the average nucleotide identity of 1,000bp fragments of crAssphage genomes from Mexico (8) and this study. **D**. Boxplots of average nucleotide identity between related mother-infant dyads sequenced in this study (Within), unrelated individuals sequenced in this study (Between), and genomes from previous studies (Global). Blue lines indicate mean average nucleotide identity. P values: NS > 0.05, * < 0.05, ** < 0.01.

**Figure 3.**
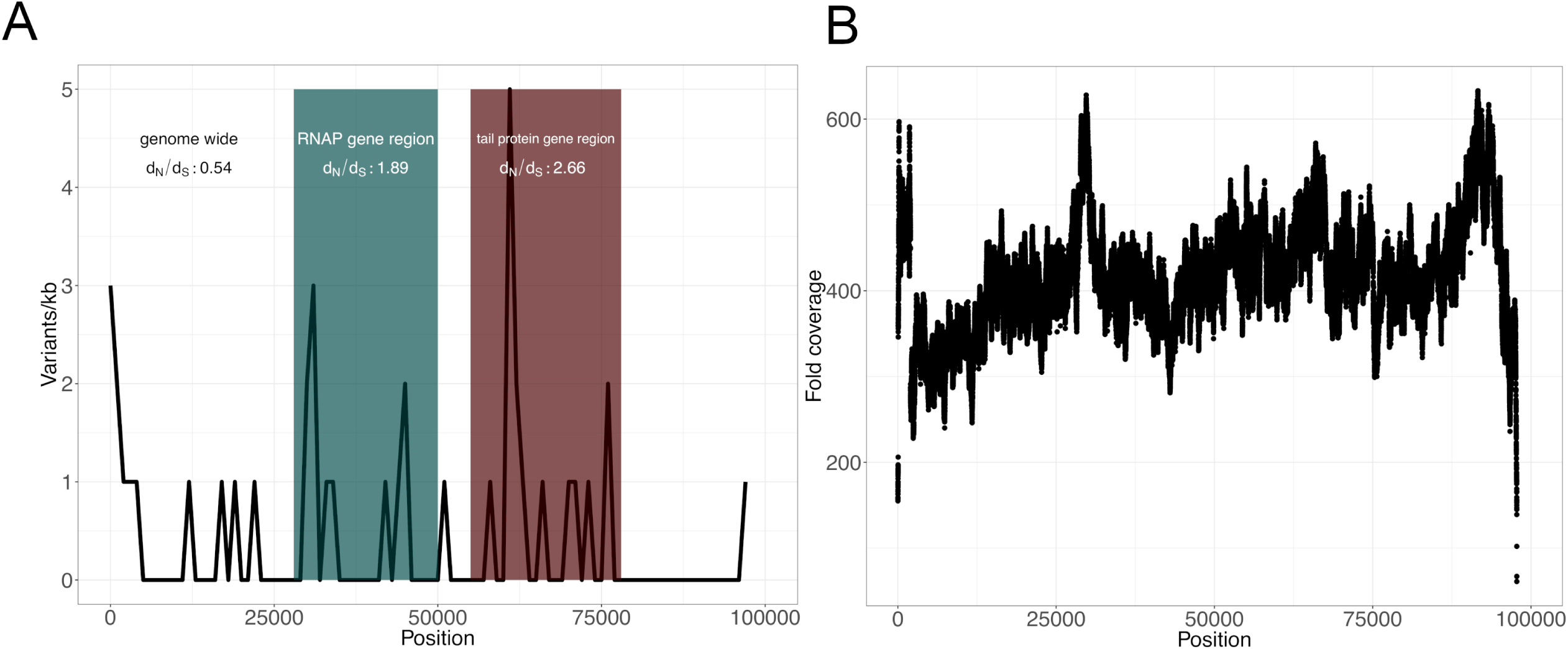
Selective pressures are variable across the genome of crAssphage M186D4. A. The distribution of variants per kilobase across the genome. The region encoding RNA polymerase genes is colored teal. The region encoding phage tail proteins is colored red. dN/dS ratios are listed in each respective region. B. The fold coverage of quality filtered reads across the genome of crAssphage M186D4.

### Average nucleotide identity and persistence across mother-infant dyads

Read mapping to the crAssphage M186D4 genome yielded complete or partially complete crAssphage genome hits in all mother-infant dyads, regardless of HIV infection or exposure status (Table 2). The percent coverage of the genome varied across samples, ranging from 2.2% to 100%. Mother-infant dyads typically yielded consistent degrees of genome coverage, though this varied across time points (Table 2). When analyzing patterns of nucleotide diversity between related maternal and infant samples, high levels of ANI between related individuals were evident (∼99%; Figure 2D), though only two mother-infant dyads had enough genomic coverage for robust assessment. Average nucleotide identities between unrelated individuals within our cohort were similar (∼97%) to levels between isolate M186D4 and previously described crAssphage variants from Mexico (Figure 2B) (8), Malawi (Figure 2C) (42), and the United States (3) (∼96%). Longitudinal sampling and analysis of infant fecal microbiota suggest that crAssphage infection persists through, at least, the first 15 weeks of life (Table 2).

**Table 2.**
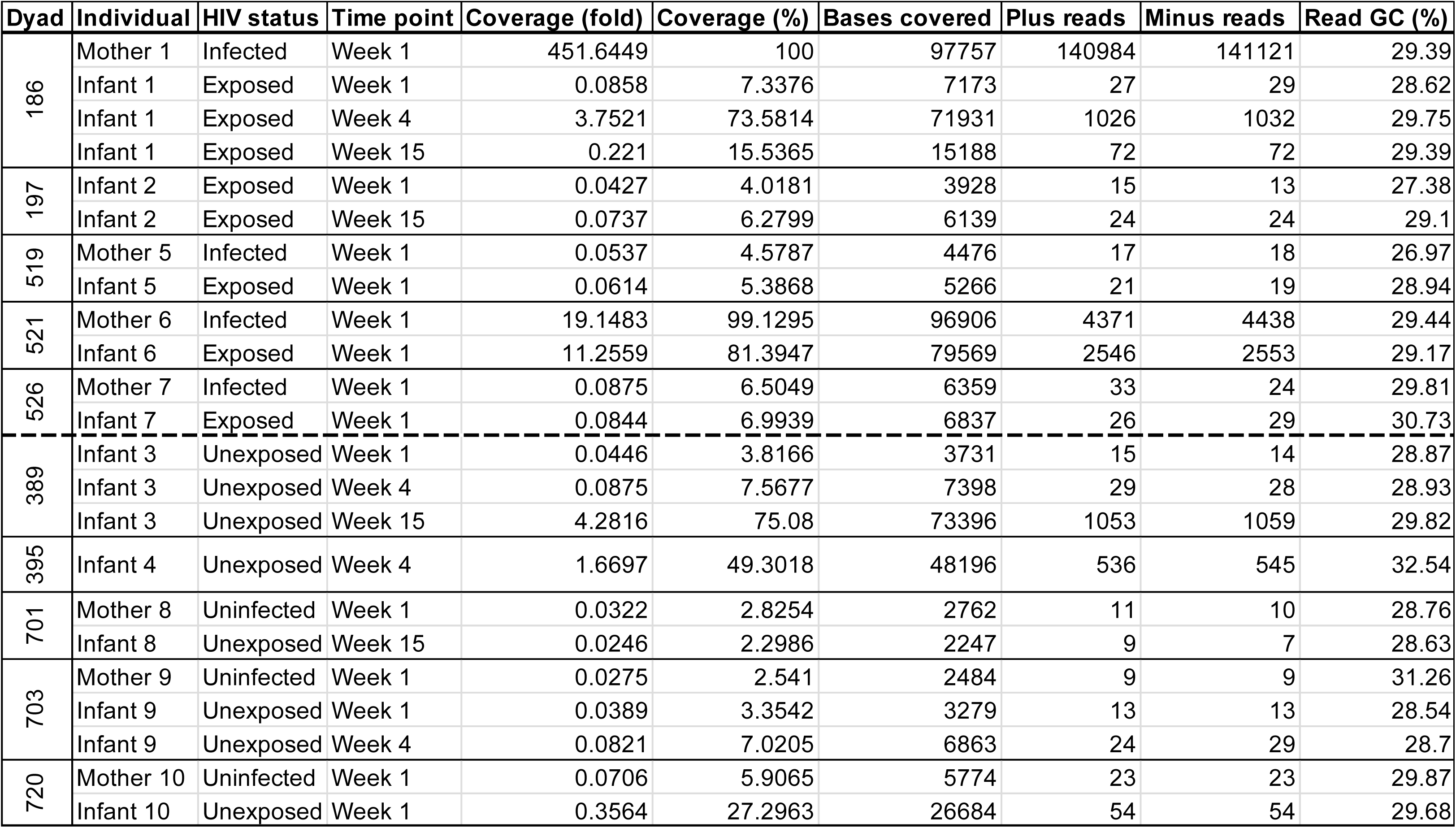
crAssphage genomic coverage across all dyads sequenced in this study. Infected/exposed dyads are separated from uninfected/unexposed by the dashed line.

### Variant analysis

We detected 40 significant single nucleotide polymorphisms (SNP) and 1 insertion/deletion in the genome of crAssphage M186D4. After filtering, 40 SNPs were retained, of which 38 fell within coding regions (Table 1), leading to a variation rate of 1 variant per 2,443 bases. The nucleotide mutation profile was composed of 33 transitions and 7 transversion, for a transition to transversion ratio of 4.71. Of the 38 variants that occurred within coding regions, 78.9% (30) resulted in nonsynonymous substitutions (*N*_*d*_), 18.4% (7) resulted in synonymous (*S*_*d*_) mutations, and only 2.6% (1) resulted in a nonsense mutation. The only insertion fell within a coding sequence for a tail tubular protein P22 (Table 1).

There was a nonuniform distribution of SNPs across the genome of M186D4. Our results show an accumulation of mutations in two distinct genomic regions (Figure 3). These mutations accumulated in RNA polymerase genes that lie within genomic regions from ∼30,000-50,000bps and phage tail protein in genomic regions from ∼58,000-76,000bps. Averaged across the genome, the d_N_/d_S_ ratio was 0.54. When considering only RNA polymerase genes, the d_N_/d_S_ ratio was 1.89. In phage tail proteins, we report a further elevated d_N_/d_S_ ratio of 2.66. The gene with the high SNP count was a putative phage tail-collar protein (DUF3751), which had a total of seven SNPs. The variant with the highest frequency across the dataset was a T -> C transition occurring within a putative Bacteroidetes-associated carbohydrate-binding (BACON) domain containing protein (Figure S1). When removing genes encoding RNA polymerase subunits and phage tail proteins from consideration, the genome-wide d_N_/d_S_ ratio fell to 0.34.

To ensure that these results were consistent across crAssphage taxa, we performed variant annotation and analysis on an additional crAssphage genome assembled from a cohort in the United States (3). As observed for crAssphage M186D4, we detected a nonuniform distribution of SNPs across the genome of crAssphage SRR073438, despite even sequencing depth (Figure S2). Consistent with observations from M186D4, SNPs persistently occurred in genes annotated as RNA polymerase and tail proteins, often resulting in nonsynonymous substitutions (Table S2). For crAssphage SRR073438, the genes with the highest SNP counts were a putative tail fiber protein and putative tail protein UGP073, both with three nonsynonymous substitutions (N_d_) and zero synonymous substitutions (S_d_, Table S2). Because all synonymous substitutions (S_d_) occurring at significant levels were distributed across protein coding genes other than RNA polymerase and tail protein genes, we were not able to calculate d_S_ values for those genes. However, average d_N_ values were similar to those observed in tail protein genes in crAssphage M186D4 (SRR073438: 0.0017, M186D4: 0.0029). Similarly, for RNA polymerase genes, d_N_ values were comparable between both genomes (SRR073438: 0.0002, M186D4: 0.0008).

### Bacterial-crAssphage interactions

We report differential shifts in bacterial abundance between samples with high and low abundance (genomic coverage) of crAssphage. We chose to use genomic coverage, rather than read count, to mitigate compositional effects and amplification bias associated with rolling circle amplification. Though genomic coverage will not eliminate bias, it remains relative to the crAssphage genome size, which appears to be comparatively static across locations and studies (9). This is in contrast to relative abundance and read counts, which are compositional in nature (43), and which are further skewed by amplification biases (44). Eleven members of the order Bacteroidales were found to have moderate or strong correlations (absolute Pearson’s *r* > 0.4) with crAssphage abundance (Figure 4a). Of these, strains of *Bacteroides thetaiotaomicron* and *Parabacteroides merdae* displayed significantly different CLR abundances between samples with high and low crAssphage abundance. *B. thetaiotaomicron* was significantly elevated in samples with high crAssphage abundance and *P. merdae* abundance was significantly reduced in samples with high crAssphage abundance. To infer interactions between the abundance of these taxa and crAssphage abundance beyond binary comparisons, we generated isometric log ratio balances between these two taxa. We report that crAssphage abundance (fold coverage) is a significant predictor (P < 0.014) of the abundance of *B. thetaiotaomicron* and *P. merdae*, independent of HIV infection/exposure status (Figure 4b).

**Figure 4.**
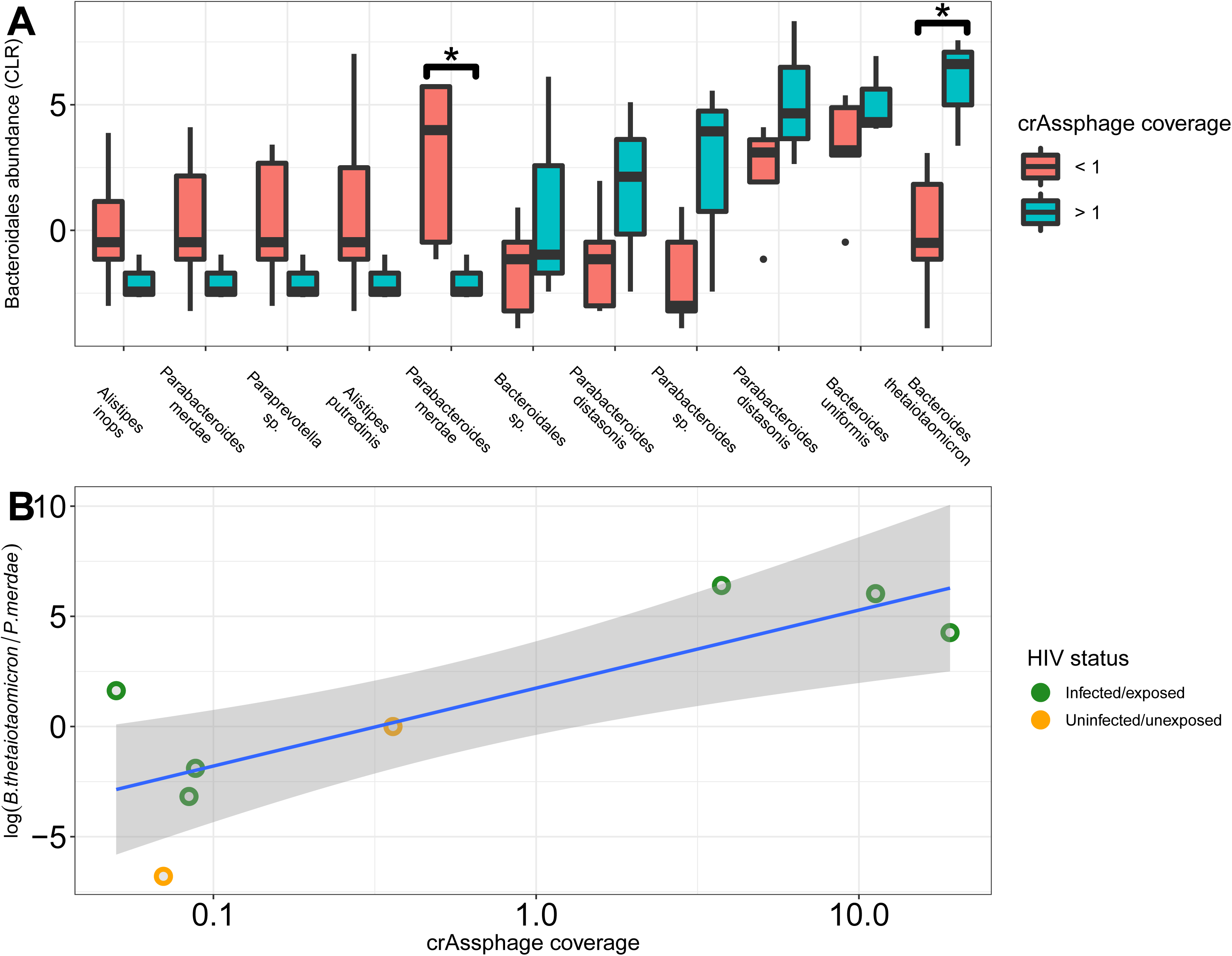
crAssphage abundance (coverage) predicts abundance of select Bacteroidales taxa. **A**. Centered log ratio (CLR) abundance of Bacteroidales taxa correlated with crAssphage abundance. CLR abundance of *P. merdae* is significantly decreased in samples with greater than 50% crAssphage coverage (red) as compared to samples with less than 50% crAssphage coverage (blue). CLR abundance of *B. thetaiotaomicron* is significantly increased in samples with greater than 50% crAssphage coverage. **B**. crAssphage coverage is a significant linear predictor of the isometric log ratio balance of *B. thetaiotaomicron* to *P. merdae*, independent of HIV infection/exposure status. HIV infected/exposed (green), HIV uninfected/unexposed (yellow), P values ≤ 0.05 are indicated with an asterisk (*).

## Discussion

As an ecological niche, the gut environment imposes a dynamic range of selective pressures on the genomes of gut microbiota. We assembled and analyzed a circularized full genome of a human crAssphage with ∼97% sequence similarity to that of the first described crAssphage genome [2]. We identified differential selective pressures along the genome, with hotspots of selection targeting groups of genes. We present evidence of substantial genetic heterogeneity and dynamic selective pressure across the genome of South African crAssphage M186D4. Many nucleotide polymorphisms occurred in protein coding regions and yielded a high ratio of substitution rates at nonsynonymous to synonymous sites (d_N_/d_S_) differentially across the genome, specifically implying strong positive selection at RNA polymerase genes and phage tail protein genes, and purifying selection genome wide. We detected complete or partially complete crAssphage genomes in all mother-infant dyads, regardless of HIV infection or exposure status. Longitudinal sampling and analysis of infant fecal microbiota suggest that crAssphage infection persists through, at least, the first 15 weeks of life. We analyzed patterns of nucleotide diversity between related maternal and infant samples, reporting high levels of average nucleotide identity between related individuals and levels of nucleotide similarity between unrelated individuals ∼2-3% lower and comparable to levels from previously published crAssphage variants from elsewhere in the world. Further, we detected differential abundance of *B. thetaiotaomicron* and *P. merdae* between samples with high and low crAssphage abundance.

The genomic selective pressures that we detected were primarily focused on two regions undergoing relatively elevated positive selection. These regions, from 30-50kb and 58-73kb, are comprised of genes from various clusters of orthologous groups (COGs), but the variation was localized into genes annotated as RNA polymerase subunits and phage tail proteins, respectively. Phage tail proteins are essential in mediating bacterial cell attachment and genome delivery. The putative protein with the highest SNP count and most variants (seven) is the phage tail fiber protein (DUF3751). This protein, while functionally uncharacterized, has been shown to mediate antagonistic interactions with bacterial taxa (45) and is a conserved element in phage-derived bacterial tailocin complexes (45). Tailocins are bacterial protein complexes co-opted from bacteriophage that are morphologically and functionally similar to phage tail proteins and are critical to eukaryotic and bacterial cell binding (46, 47). Furthermore, T4 tail adhesion protein, which is homologous to DUF3751, mediates adsorption of T4-like bacteriophages to *Escherichia coli* and contains a lipopolysaccharide (LPS) binding site. T4 tail protein gp12 has been shown to modulate host inflammatory responses to LPS *in vivo* (48).

Though it is unclear how this domain physically interacts with bacterial cell receptors, positive selection along this gene and other tail proteins may represent a Red Queen scenario where the phage tail protein is adapting to constantly evolving target cell surface proteins. Adaptive evolution of phage tail proteins has been well documented in marine bacteriophage communities (49) and has been shown to facilitate an expanded host range. Additionally, long-term (23 day) phage/*B. intestinalis* co-cultivation experiments have shown that founder strains of crAssphage have severely limited ability to infect bacterial strains passaged for the length of the experiment (11), indicating rapid, likely antagonistic, coevolution between crAssphage and bacterial host strains. This reduction in infection rate likely reflects nonsynonymous changes in bacterial cell surface receptors that are mediated by interaction with crAssphage tail proteins.

We detected genomic regions of crAssphages in all mother-infant pairs and samples sequenced, regardless of HIV infection/exposure status or time point sampled (Table 2). Interestingly, the percent of genome coverage from read alignment to isolate M186D4 was relatively consistent within a mother-infant pair. Though this observation is not a truly quantitative assay of viral abundance, consistent coverage of the crAssphage genome between related mother and infant samples at one week postpartum is suggestive of comparable abundance of crAssphage taxa, potentially from vertical transmission. Importantly, this differs from findings in infants from Missouri, USA, where crAssphage-like sequences were only identified at 24 months and not in earlier samples (50). We identify crAssphage sequences in feces as early as 4 days of age. We also note that genomic coverage of crAssphage in the same infant varied across time, with differential coverage during later sample points.

At the nucleotide level, our results support vertical transmission of crAssphage from mothers to infants. We report elevated average nucleotide identities (∼99%) shared between genomes from related mothers and infants, as compared to unrelated pairs (∼97%). This trend was consistent between maternal and infant samples collected at 4 days postpartum, but also held true for infant samples collected at week 4. Whether this is due to continued viral transmission via breastfeeding or other sources remains unclear. Average nucleotide identities between unrelated individuals within our cohort were similar to, though slightly higher than, levels between the genomes of isolate M186D4 and the three previously described global strains with complete genome sequences. Overall, our data are suggestive of maternal transmission of crAssphage to infants, though further investigation is needed to accurately quantitate viral load and patterns of nucleotide diversity between mothers and infants, as well as over longitudinally collected samples starting at birth to assay viral dynamics.

With regard to the gut bacterial cohort, we report altered bacterial dynamics between samples with high and low crAssphage abundance. Previous studies have demonstrated that members of the Bacteroidales are preyed upon by crAssphage taxa (2, 11), though the dynamics of this relationship are unclear. In this cohort, we report significantly altered abundance of *B. thetaiotaomicron* and *P. merdae* in samples with high versus low crAssphage abundance, independent of HIV infection/exposure status. Using compositional data analytical approaches (isometric log ratio balances), we report that crAssphage abundance is a significant predictor of the log ratio abundance of these taxa (Figure 4b). *B. thetaiotaomicron* has been previously hypothesized as a potential host of crAssphage taxa (2, 9), though our data are the first to demonstrate this positive correlation. *P. merdae* belongs to the recently reassigned genus, *Parabacteroides*, which is sister to *Bacteroides* and a persistent member of the gastrointestinal tract (51, 52). The opposite shifts in abundance between these two taxa may represent unique relationships with crAssphage taxa (e.g. non-disruptive proliferation in *B. thetaiotaomicron* as reported for *B. intestinalis* (11), or comparably elevated lytic activity in *P. merdae*) or shared ecological niches in the gastrointestinal tract. In scenarios where crAssphage and *B. thetaiotaomicron* are elevated, *P. merdae* may decrease in abundance due to lack of resource access or other antagonistic bacterial interactions, though further studies dissecting this relationship are needed.

In summary, our data argue for vertical transmission of crAssphage across mother-infant dyads, potentially as a component of the inherited microbiome. Our results suggest genome-wide purifying selection in crAssphage, with episodic shifts of strong positive selection within RNA polymerase and phage tail protein genes. Elevated selective pressure of phage tail proteins may be due to antagonistic coevolution between crAssphage and bacterial targets, though further work is needed to demonstrate this conclusively. Additionally, our data suggest that crAssphage abundance may direct gut bacterial dynamics, providing an ecological basis for the associated genomic consequences. Collectively, our results argue that crAssphage alters bacterial consortia and may act as a driver of ecological and evolutionary dynamics in the gut.

## Acknowledgements

This study was funded in part by the University of Washington Center for AIDS Research, an NIH funded program under award number AI027757, supported by the following NIH Institutes and Centers (NIAID, NCI, NIMH, NIDA, NICHD, NHLBI, NIA, NIGMS, NIDDK). The InFANT study cohort was supported in part by the Canadian Institutes of Health Research HIV Vaccine Initiative grant (#01044-000), NIH R01AI131302 and AI120714-01A1. We would like to thank the InFANT study team for collecting samples. We also thank all participants in the study for providing samples. DC is supported by a Wellcome Trust DELTAS Africa grant to the SANTHE programme (grant #107752/Z/15/Z).

## Contributions

HBJ, DC, AV and DM designed the study. JW and DDN prepared bacterial 16S rDNA libraries. DC, EH, SJ prepared the samples for sequencing. BPB and AV analyzed the data. HBJ, BPB, and AV wrote the manuscript. All authors reviewed and edited the manuscript.

---

Figure S1. Variant frequency across the genome of crAssphage isolate M186D4. Each significant variant is indicated by a point. Lines are added for visual aid only.

Figure S2. The fold coverage of quality filtered reads across the genome of crAssphage SRR073438.

Table S1. Single nucleotide variants across the genome of crAssphage M186D4.

Table S2. Functional annotation of variants in crAssphage SRR073438

